# Does the Freeze-all strategy improve the cumulative live birth rate and the time to become pregnant in IVF cycles?

**DOI:** 10.1101/2020.06.10.144055

**Authors:** S. Johnson, J. Vandromme, A. Larbuisson, D. Raick, A. Delvigne

## Abstract

**Introduction:** Freezing of all good quality embryos and their transfer in subsequent cycles, named the freeze-all strategy (FAS), is widely used for ovarian hyperstimulation syndrome (OHSS) prevention. Indeed, it increases live birth rates among high responders and prevents preterm birth and small for gestational age. Consequently, why shouldn’t we extend it to all?

**Materials and methods:** A retrospective and monocentric study was conducted between January 2008 and January 2018 comparing the cumulative live birth rates (CLBR) between patients having undergone FAS and a control group using fresh embryo transfer (FET) and having at least one frozen embryo available. Analyses were made for the entire cohort (population 1) and for different subgroups according to confounding factors selected by a logistic regression (population 3), and to the BELRAP (Belgian Register for Assisted Procreation) criteria (population 2).

**Results:** 2216 patients were divided into two groups: Freeze all (FA), 233 patients and control (C), 1983 patients. The CLBR was 50.2% vs 58.1% P=0.021 for population 1 and 53.2% vs 63.3% P=0.023 for population 2, including 124 cases and 1241 controls. The CLBR stayed in favour of the C group: 70.1% vs 55.9% P=0.03 even when confounding variables were excluded (FA and C group respectively 109 and 770 patients). The median time to become pregnant was equally in favour of the C group with a median of 5 days against 61 days.

**Conclusions:** CLBR is significantly lower in the FA group compared to the C group with a longer time to become pregnant. Nevertheless, the CLBR in the FA group remains excellent and superior to that observed in previous studies with similar procedures and population. These results confirm the high efficiency of FAS but underline the necessity to restrict the strategy to selected cases.

## Introduction

Freezing all good quality embryos and their transfer in subsequent cycles, named as the freeze-all strategy (FAS), is widely used for ovarian hyperstimulation syndrome (OHSS) prevention. This technique has the advantage of transferring the embryos into a more physiological environment (Roque et al., 2015). Indeed, transferring frozen-thawed embryos during subsequent cycles avoids the hyper-oestrogenic climate that results from ovarian stimulation, which has a negative effect upon synchronisation of the embryo and the endometrium (Roque et al., 2019; Perrier d’Hauterive et al., 2002). Embryo implantation is a crucial stage in IVF techniques and implantation failure remains difficult to resolve. Endometrial receptivity seems to be implicated in 66% of the cases as only 33% is due to the embryo itself (Achache H et al., 2006). This is why, in addition to the benefit of the FAS in OHSS prevention, several studies have shown the benefits of FAS on live birth rates among high responders (Evans et al., 2014; Roque et al., 2013; Chen et al., 2016, Bosdou et al, 2018). Moreover, the benefits of frozen embryo transfer (FRET) compared to fresh embryo transfer (FET) have been demonstrated to prevent preterm birth and small for gestational age (Maheshwari et al., 2018). Consequently, why should we limit the FAS to high responders rather than extend it to all? Previous studies were conducted but the results were inconsistent. A meta-analysis in 2018 included 11 studies with over 5000 patients (Roque et al., 2018). The live birth rate was in favour of the freeze all group compared to the group with FET (46% vs 42.7% P=0.04), but the patient subgroups revealed that this was true only for hyper-responders. A prospective randomised trial, included in this meta-analysis, conducted over 2157 patients compared the live birth rate between patients without any ovulatory disorder who had a fresh versus frozen embryo transfer (Shi et al.,2018). No difference was found between the two groups. This observation was not confirmed in another large prospective trial of 1650 normo-ovulatory patients (Wei, 2019). Finally, the only retrospective study which analysed the cumulative live birth rate after the first IVF cycle using the freeze all strategy found a CLBR of 50.74% but had no control group. With these conflicting results, no conclusions could be made. We therefore conducted a new real-life study on FAS in which we focused on the CLBR in our overall IVF population. We compared an FA group in which “non-elective” freezing was applied (Blockeel et al.,2019) with a control group which, to be compared, also had frozen embryos available to be transferred following the fresh ones.

## Materials and Methods

This is a retrospective and monocentric study conducted between January 2008 and January 2018 comparing the CLBR and the time taken to become pregnant in patients having undergone FAS to those using FET and having at least one frozen embryo ready to be transferred.

### Ethical approval

The study protocol was approved by the ethics committee of the Saint-Vincent Clinic CHC on the 10^th^ April 2019 n° 100409.

### Primary outcomes

The first primary outcome was the cumulative life birth rate. It was defined as the rate of live births for one patient after one OPU until the first live birth (following the embryo transfer (fresh or frozen) or after having used all the frozen embryos of this OPU. A live birth was defined as an infant born alive after 24 weeks of gestation

The second primary outcome is the time taken to become pregnant defined as the time between the OPU and the transfer that results in the live birth.

### Secondary outcomes

Secondary outcomes were: Clinical pregnancy rate, miscarriage rate, cumulative live birth rate according to the number of oocytes retrieved. Clinical pregnancy was defined as an ongoing pregnancy determined by ultrasonographic documentation with at least one foetus with a heartbeat. Miscarriage was defined as a pregnancy loss before 22 weeks of gestation. The number of oocytes retrieved were calculated for one OPU and classified in categories: 1-5, 6-10, 11-15, 16-20, 20-25, ≥ 26.

### Population

The data were obtained from the IVF database of ART centre CHC Montlégia. They were extracted by a statistical request in an independent and anonymous file. Women who underwent an IVF cycle between January 2008 and December 2018 were selected. One cycle per women was selected. The study analysed two groups: the “freeze all” (FA) group and the control (C) group:

In the FA group, all embryos obtained after oocyte retrieval were cryopreserved and transferred in subsequent cycles.

The C group represents women who had a fresh transfer following oocyte retrieval, with at least one frozen embryo during their IVF cycle.

Analyses were made for the entire cohort (population 1) and also for different subgroups according to confounding factors selected by the Logistic Regression (LR) (population 3) and to the BELRAP (Belgian Register for Assisted Procreation) criteria used for our national benchmark (population 2).

Belrap criteria are used to construct the annual benchmark for all Belgian centres considering patients less than 36 years old, undergoing their first IVF trial.

To calculate the CLBR, we limited the gathering of information to January 2018 in order to concentrate on birth related statistics (9 months + information retrieval). The women who were not pregnant at the time of the study but who had frozen embryos left were excluded for the first end point.

### Ovarian stimulation

Women were monitored and managed according to our institutional clinical protocols. Various protocols were applied using between 100 and 300 IU of gonadotropins per day for ovarian stimulation according to the patient’s age, body mass index (BMI) and Anti Mullerian hormone (AMH) rate. Recombinant Follicle-Stimulating hormone (FSH) (Puregon-MSD, Courbevoie, France; Gonal-F, Merck, Genève, Suisse) or urinary FSH (Menopur, Ferring pharmaceuticals, Alost, Belgium) was used in a GnRH antagonist protocol, a long agonist protocol or a short agonist protocol. Pituitary down-regulation was obtained with a GnRH antagonist (Cetrotide, Merck) or a GnRH agonist (Busereline, Suprefact, Sanifi-Avantis, Diegem, Belgium; Gonapeptyl, Ferring). Final Oocyte maturation was triggered when ≥3 ovarian follicles of ≥ 17 mm were visible by ultrasound. Final Oocyte maturation was achieved using either a single injection of 0.2mg of GnRH agonist (Tryptoreline, Decapeptyl, Ipsen, Signes, France) or 5000 IU of human Chorionic Gonadotropin (hCG) (Pregnyl, MSD, Brussels, Belgium) according to the OHSS risk factors (Dosouto C et al., 2017).

### Oocyte insemination

Conventional IVF or ICSI were performed according to sperm parameters. In conventional IVF, oocytes were inseminated during the night with a sperm suspension containing 100 to 150×10^3^ mobile spermatozoid/ml. With the ICSI method, the cumulus was destroyed using an hyaluronidase at the concentration of 40 IU/ml. The sperm injection was executed during metaphase two, four hours after OPU. Fertilization was assessed by the presence of two pronuclei (2PN) 16 to 18 hours after oocyte insemination.

### Embryo culture, cryopreservation, and thawing

Embryos were cultured at 37 °C in incubators with 6% of CO2 and 5% of O2. Each day, embryo morphology was evaluated and classified according to Istanbul Criteria (Balaban B et al., 2001). On day 3, we considered whether all embryos should be cultured until day 5 rather than transferring the best at day 3. If on day three, 4 or more embryos had a perfect morphological score (score 8-cell, grade 1 or 7-cell, grade 1) the entire cohort was cultured until day 5. The best quality embryo was then chosen for transfer. If 2 embryos had the same score, the one with the best kinetic development was transferred. Each additional embryo, or embryos selected for freeze-all, were frozen. The freezing technique changed in January 2011 from slow freezing to vitrification.

Transfer policy in our IVF centre: Belgian law only allows transfer of a single fresh embryo in women under 36 years old undergoing their first trial. Nevertheless, the transfer of two frozen-thawed embryos is allowed for such patients. The combination of Belgian law and our methods of embryonic culture (vitrification of blastocysts since 2011) has led to unacceptable rates of multiple pregnancies when transferring two frozen-thawed embryos. For this reason, we currently encourage the transfer of a single selected blastocyst in patients under 36 years old (Raick D et al., 2014).

Internal practise to avoid OHSS was to always perform FA when 25 oocytes or more were retrieved. Between 15 and 25 the decision was left up to the gynaecologist who performed the OPU.

FA policy was also applied for:

- Uterine anomalies detected during ovarian stimulation (de Ziegler et al. 2016; Tsiami et al. 2016; liu et al, 2018) which includes endometrial, myometrial, or tubal anomalies
- progesterone levels superior to 1.5ng/ml on the day of ovulation triggering (Lawrenz et al.,2017; Asada Y. et al, 2019; Vuong L. N. et al., 2019).

### Data analysis and statistics

The general characteristics of the patients in both groups were recorded prospectively. The following data were collected: age (in years), BMI, type of infertility (primary or secondary), factors of infertility (idiopathic, mechanical, endocrine, male, ovarian failure) and tobacco intake. We also recorded the number of oocytes retrieved, the number of mature oocytes inseminated, the embryo development stage, the number of embryos transferred and their quality, the number of embryos left after pregnancy was obtained and the total number of transfers performed for one OPU and OHSS.

A power analysis was realised based on the results obtained for the main outcome in each population.

The statistical analysis was realised using SPSS. We used Chi-square, Kruskal-Walis or mann- Whitney and Wilcoxon testing. A p value < 0.05 was considered as statistically significant.

Logistic regression (LR) was used to identify confounding variables.

## Results

### Characteristics of the study population

The entire cohort included 2628 women; 2325 in the C group and 303 in the FA group. The different reasons for performing FA are detailed in table 1.

**Table 1:**
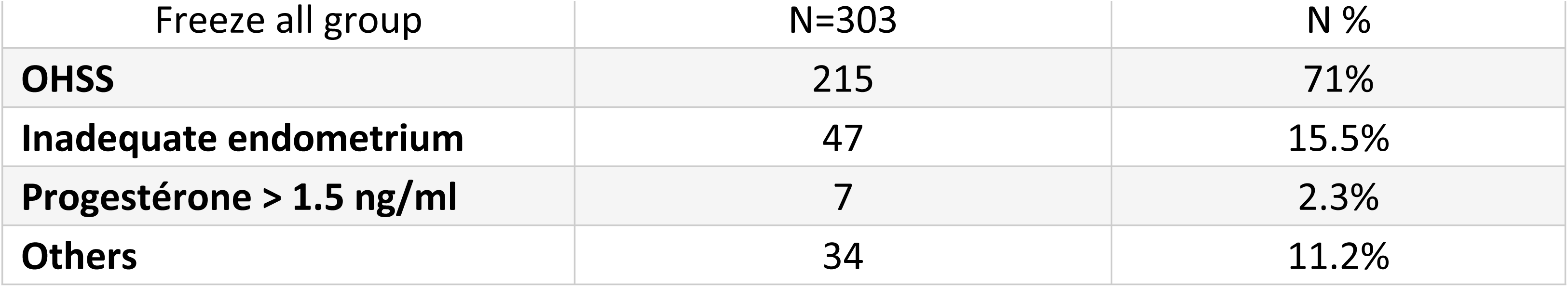
Indication to perform Freeze all.

| Freeze all group | N=303 | N % |
| --- | --- | --- |
| **OHSS** | 215 | 71% |
| **Inadequate endometrium** | 47 | 15.5% |
| **Progestérone > 1.5 ng/ml** | 7 | 2.3% |
| **Others** | 34 | 11.2% |

Women included in the FA group were mainly selected due to a risk of OHSS (71%). The other reasons FA was performed are as follows: 15.5% of the women had an inadequate endometrium, polyps or liquid in the uterus cavity and high progesterone levels before ovulation triggering (2.3%). Other lesser reasons were: patient sicknesses (HTA, infectious seroconversion), post retrieval complications, psycho-sociological problems, failed fresh embryo(s) transfer, hydrosalpinx visualised during ovarian stimulation.

### Primary outcomes

#### Cumulative live birth rate after the first IVF cycle

After excluding women who were not pregnant at the time of the study but who still had frozen embryos left, the eligible cohort included 233 women in the FA group and 1983 patients in the Control group (population 1).

A second population selection was done according to the Belrap criteria, meaning women who were undergoing their first IVF cycle and who were under 36 years of age. This second eligible cohort consisted of 124 women in the FA group, and 1241 women in the C group (population 2) (Fig 1).

**Figure 1.**
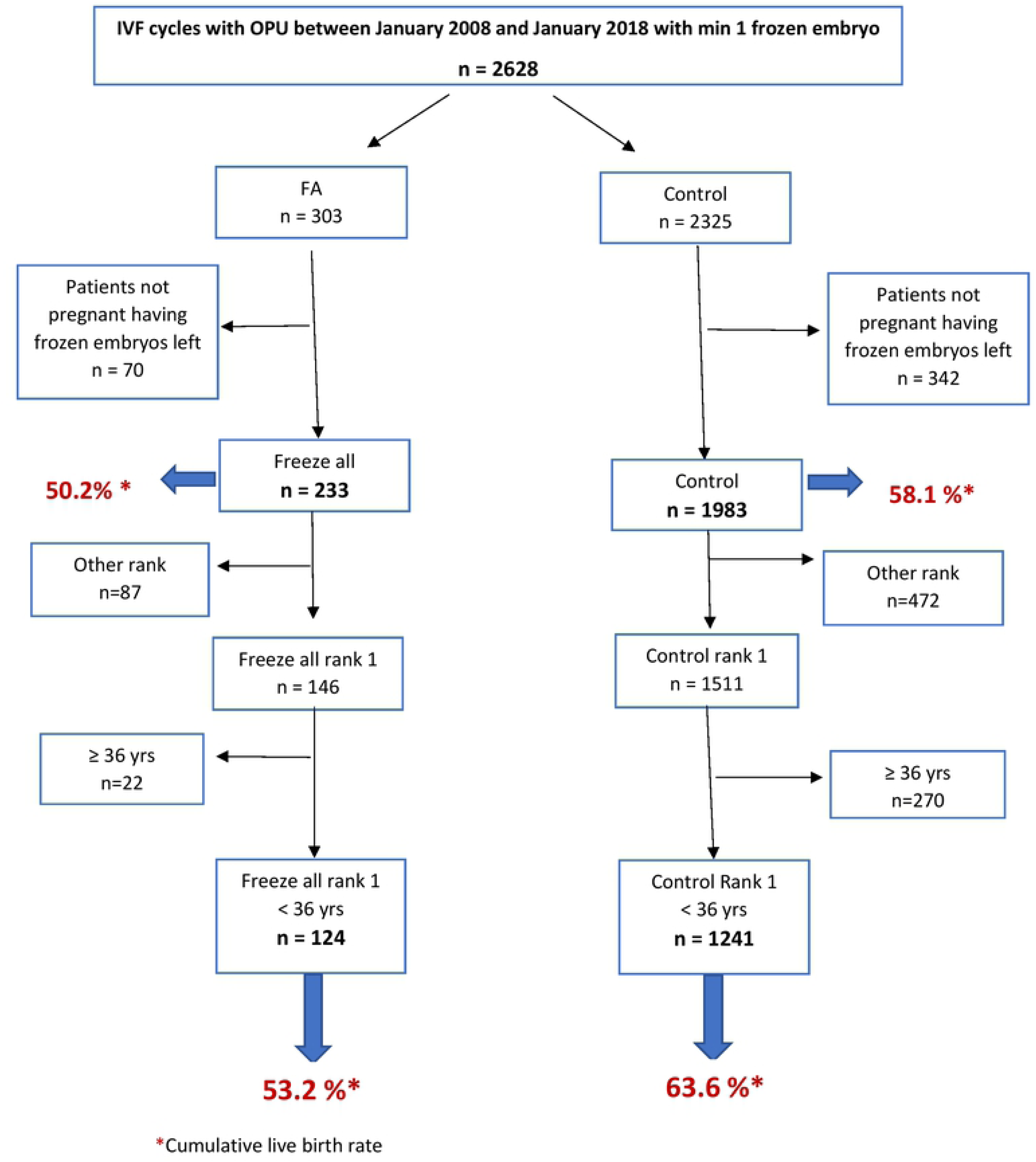

A logistic regression analysis was made to exclude the possible confounding variables. Table 2 shows the results of the LR with the following confounding variables: the patient’s age, the rank of the trial, tobacco use, the number of embryos obtained, the total number of transfers and the date of the OPU. A third analysis was made after suppression of the confounding factors found by the LR. This group included 109 patients in the FA group and 770 in the C group (population 3) (**Fig 2)**.

**Table 2:** Logistic regression and confounding factors.

|  | OR | IC 95% | P value |
| --- | --- | --- | --- |
| **Group** | 2.247 | 1.532-3.295 | **0.000** |
| **Age (years)** | 0.905 | 0.879-0.931 | **0.000** |
| **Rank** | 0.726 | 0.541-0.973 | **0.032** |
| **Primary/secondary inf. patient** | 1.249 | 0.941-1.658 | 0.123 |
| **Primary/secondary inf. partner** | 1.178 | 0.892-1.555 | 0.248 |
| **BMI (Kg/m<sup>2</sup>)** | 0.998 | 0.972-1.024 | 0.863 |
| **Tabacco patient** | 0.695 | 0.513-0.941 | **0.019** |
| **Tabacco partner** | 1.422 | 1.087-1.861 | **0.010** |
| **Infertility diag. patient** | 1.072 | 0.943-1.217 | 0.287 |
| **Infertility diag. partner** | 0.969 | 0.781-1.203 | 0.778 |
| **Number of oocytes retrieved** | 0.990 | 0.954-1.026 | 0.573 |
| **Number of obtained embryos** | 1.229 | 1.157-1.306 | **0.000** |
| **Total number of transfers** | 0.483 | 0.425-0.549 | **0.000** |
| **Mean number of transfers** | 1.127 | 0.811-1.565 | 0.477 |
| **Date of OPU after 2011** | 1.915 | 1.461-2.510 | **0.000** |

**Figure 2.**
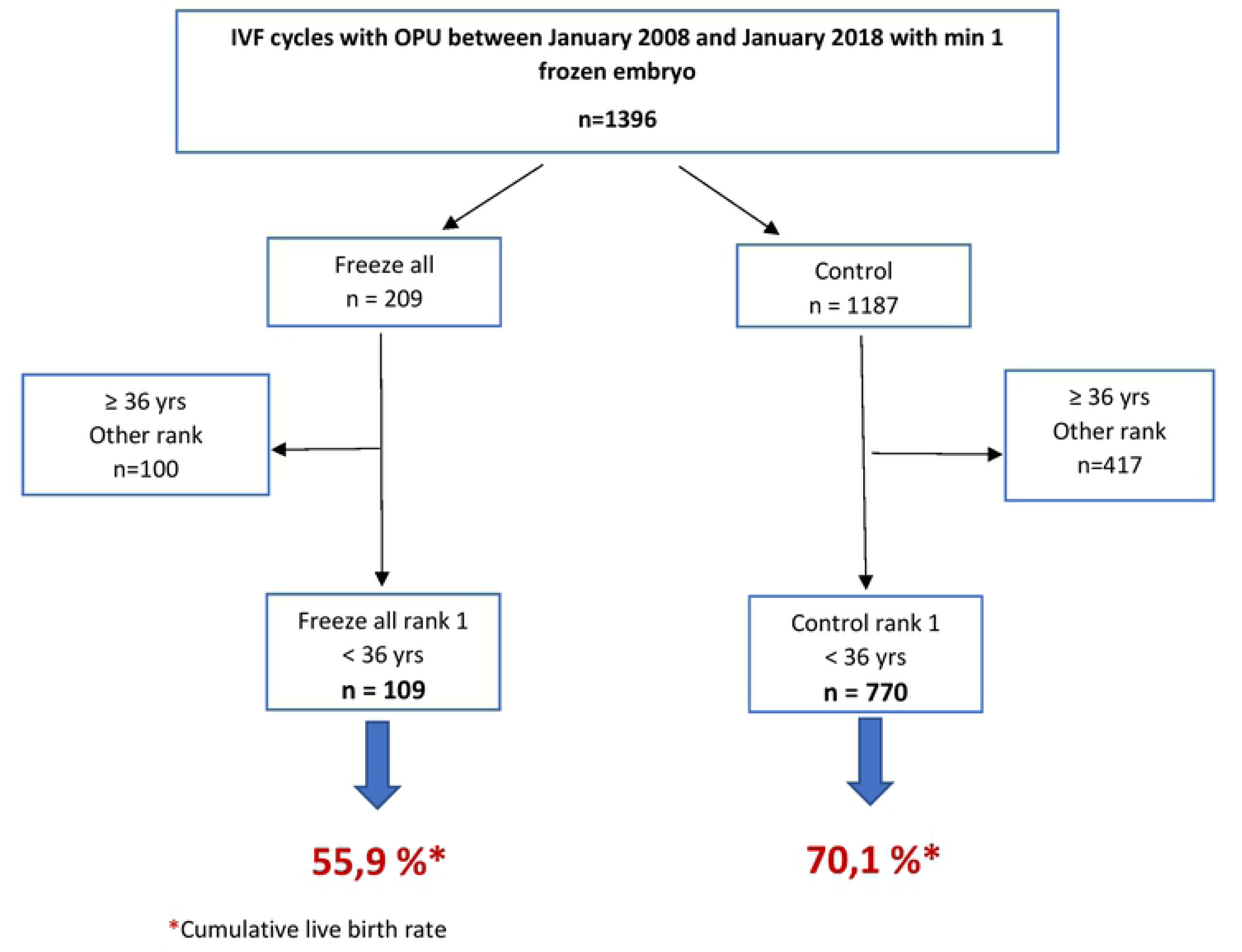

For population 1, the CLBR is 50.2% for the FA group against 58.1% for the C group (P=0.021). For population 2, the CLBR is 53.2% for the FA group against 63.6% for the C group (P=0.023). For population 3, the CLBR is 55.9% for the FA group against 70.1% for the C group (P=0.03) **(Fig 1 and 2)**.

### Time to become pregnant

The time to become pregnant is significantly shorter in the C group compared to the FA group with a median of 5 days against 61 days. **(Fig. 3)**

**Figure 3.**
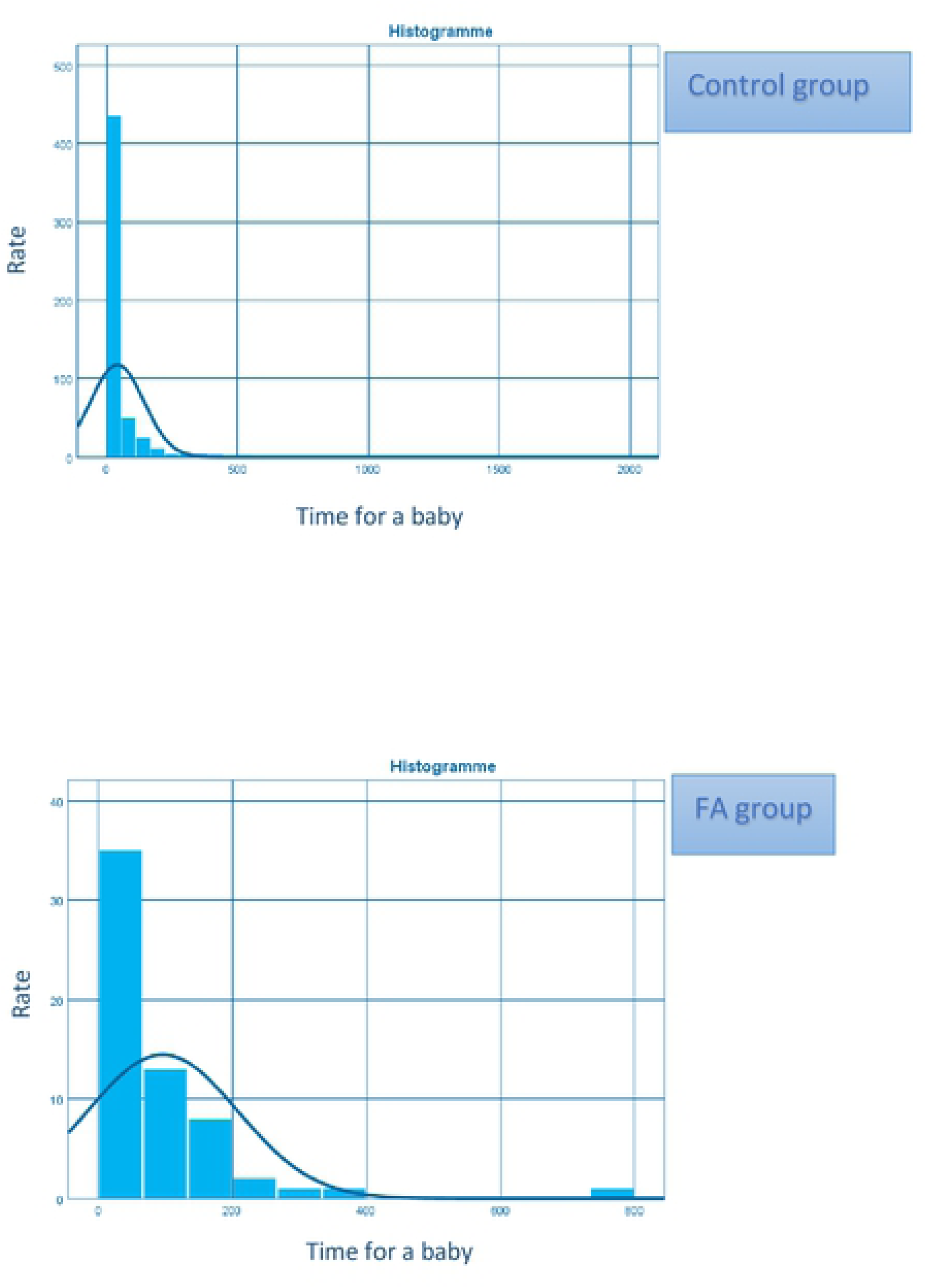

### Secondary outcomes

The results for the second end points were based on population 3 (after exclusion of confounding variables). The cumulative clinical pregnancy rate is higher in the C group with 71% against 58 % for the FA group. The cumulative miscarriage rate is similar in both groups (12.2% in the FA group and 12.7% in the C group, P=0.5).

We calculated the cumulative live birth rate according to the oocyte number as shown in figure 4. The CLBR rose from 20% to 83.33% when 21 to 25 oocytes were retrieved in the FA group. In the C group the CLBR rose from 50.13% to 68.35% when 16 to 20 oocytes were retrieved.

**Figure 4:**
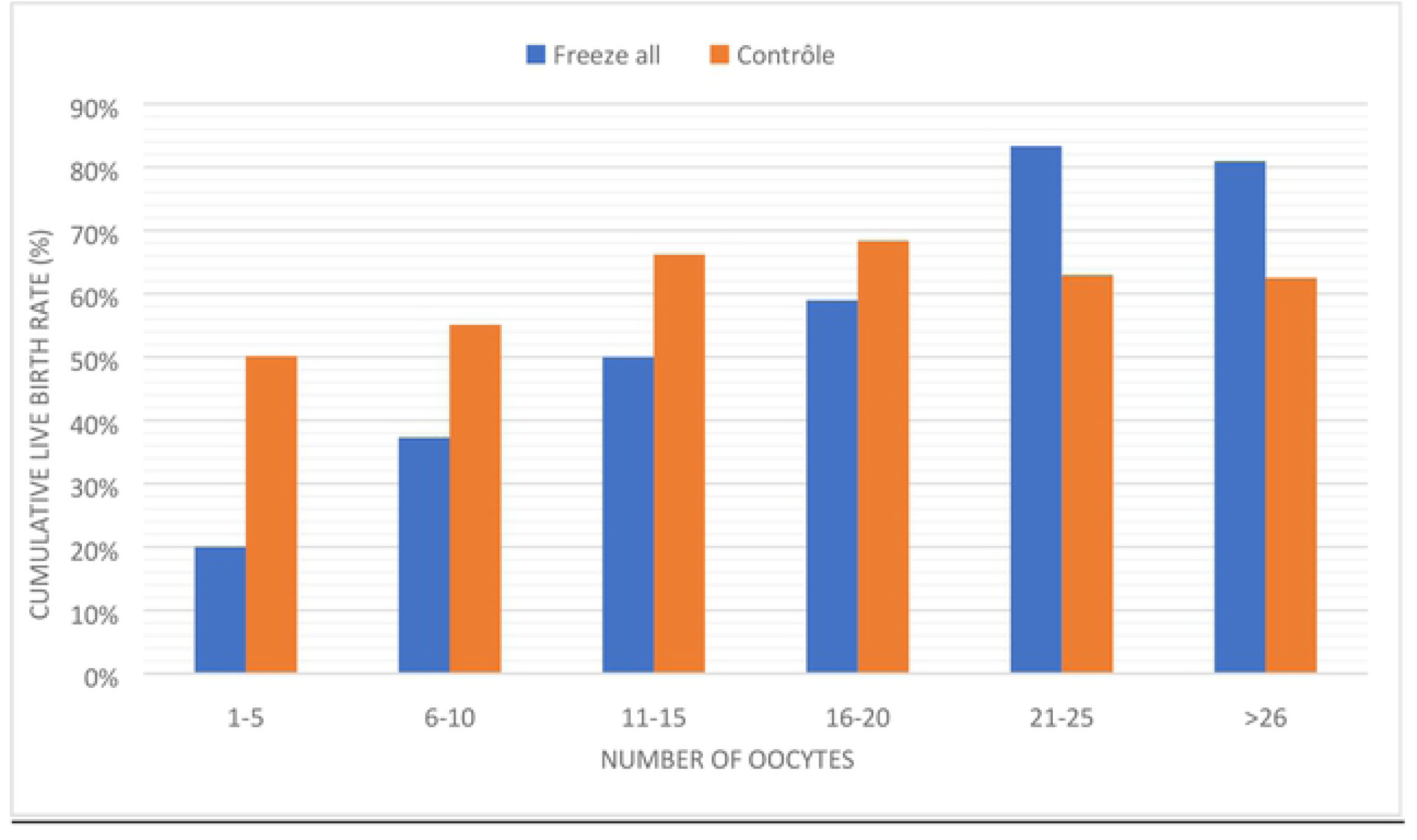
CLBR according to the number of oocytes retrieved.

### Power Analysis

A power analysis was realised based on the results obtained for the main outcome in each population.

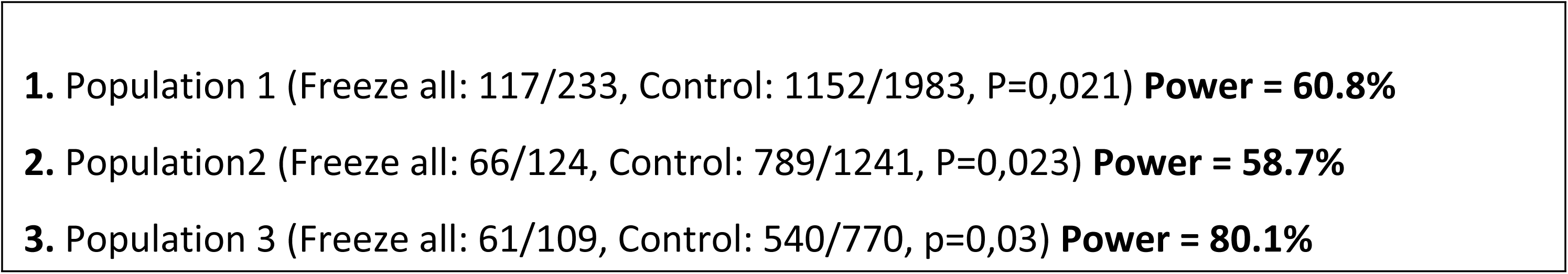

## Discussion

The advantages of the FA strategy need no further corroboration for women at risk of OHSS. (Chen et al.,2016; Shapiro et al.,2011; Aflatoonian et al., 2010). However, results remain contradictory when applying the FA strategy to a normo-responsive population (Roque et al., 2018; Shi et al.,2018, Wei et al., 2019).

This study has analysed retrospectively a cohort of patients having undergone an FA strategy compared to a control group having benefitted from a fresh transfer with remaining frozen embryos, in a “real life” study situation, the first end point being the cumulative live birth rate.

Our results show that the CLBR is significantly higher in the control group compared to the FA group (70.1% against 55.9%, P=0.03) taking into account the identified confounding factors with a power of 80%, which is sufficient.

These results contradict certain data from the most recent literature on the topic. Indeed, Shi et al. in 2018 carried out a prospective randomised trial including 2157 women without any ovulatory problem. They found no difference in live birth rates between the FA and C groups. (48.7% vs 50.2%). But only the live birth rate after the first transfer was calculated and not the cumulative live birth rate.

Roque et al. in 2018 published a meta-analysis which found a significant increase in the live birth rate in the FA group compared to the women having undergone a fresh embryo transfer in the overall IVF population (46% against 42.7%). But no difference was observed when the cumulative life birth rate was considered among the same population (RR = 1.04; 95% CI: 0.97–1.11).

The efficiency of freeze-all strategy calculated by cumulative pregnancy rate (and not birth rate) is underlined by Zhu et al. in his retrospective study of 20687 patients published in 2018. His results show a cumulative pregnancy rate of 50.74% but this study does not benefit from control group results. Finally, a multicentric randomised controlled trial showed that frozen single blastocyst transfer resulted in a higher singleton live birth rate than did fresh single blastocyst transfer in ovulatory women (Wei et al., 2019).

Even though, some other recent studies corroborate our results (Smith A.D.A.C. et al., 2019). In our study, the performances of the freeze all strategy can also be underlined with a CLBR of 50.2% for the overall population and 55.9% in population 3 (after exclusion of the confounding variables). Even if these results are lower than the one in the Control group, a CLBR of 55.9% is the best rate published for FA in a population of women under 36. Notably, a CLBR of 93.82% was found in a recent study (Zhao et al., 2020) but it concerns a very young sub-group of women with an average age of 28. We must underline that the CLBR in the C group (70.1%) is particularly high, and so significantly better than the FA group. These particularly good results can probably be explained by a selection bias in our control group, which only concerns patients with fresh and frozen embryos for this first IVF trial. Why these are such “good patients” may be explained by superior “receptiveness”, as their oocyte quantity and quality allowed at least one frozen embryo, in addition to the fresh embryo(s) transferred. This relationship between the number of embryos cryopreserved after a fresh transfer and the CLBR was confirmed by a recent work. (Scaravelli et al.,2019).

Less efficient vitrification techniques could explain the differences between our and previously published results in favour of FA. This seems unlikely when considering the CLBR in the FA group reflecting good quality thawed embryos. Moreover, the pregnancy rates after frozen-thawed embryo transfer in our study during the same period is 35.4% for clinical pregnancies and 27.6% for ongoing pregnancies (excluding miscarriages and extra- uterine pregnancies). Our vitrification procedure seems therefore not in question (Papanikolaou E. et al., 2019). Nevertheless, we must underline that one confounding factor is the date of OPU, before or after 2011, which is when our technique of cryopreservation changed from slow freezing to vitrification.

The time taken to obtain a first pregnancy is statistically and clinically significant in favour of the Control group. (5 times longer for the FA group). These results attest also to the good performance obtained from the first fresh embryo transfer among a particularly receptive group. We always try to reduce this time to the minimum by transferring the frozen embryos as soon as feasible during the following cycle. The first thawed embryo transfer can take place at least 1 month after oocyte retrieval. Indeed, a study published in 2018 (Bourdon et al., 2018) demonstrated that pregnancy rates did not decrease if the transfer of the frozen-thawed embryo was made the cycle immediately following the OPU. The large difference in the time taken to become pregnant may also be explained by a high proportion of high responders among the FA group, who required more time for inclusion in a new treatment cycle, needing to wait for ovarian and pelvic returns to normality. By comparison, another retrospective study found a median time of 38 days for the FA, there being no control group (Ozgur et al., 2019). This difference is difficult to explain, and highlights the many variables at work, such as the medico-social environment in each country, the accessibility to ART, and internal medical protocols.

To explain the difference in CLBR between groups, the mean number of transferred embryos per transfer was analysed (data not communicated). Belgian law only allows the transfer of a single fresh embryo but does allow the transfer of two frozen-thawed embryos for women under 36 years old undergoing their first try. However, in this group (population 2), the number of transferred embryos was not statistically different between groups (1.26 for FA and 1.23 for control group (P=0.37).

We have also analysed the cumulative live birth rates according to the number of oocytes retrieved. In recent years, many studies have attempted to define the relationship between these two IVF data. Contradictory results were observed in three retrospective studies evaluating the live birth rate (Sunkara et al., 2011; Cai et al., 2013; Steward RG et al., 2014). Two more recent studies have considered CLBR in relation to the number of oocytes obtained. Drakopoulos et al. analysed 1099 IVF cycles for CLBR using fresh and frozen-thawed embryo transfers. No relationship was found between LBR and the number of oocytes retrieved when considering only fresh embryo transfers. However, the CLBR does increase with the number of oocytes retrieved when both fresh and frozen-thawed embryo transfers are taken into account. (Drakopoulos et al., 2016). Another retrospective study collected data from 15 IVF centres for the same analyses and observed a significant increase in CLBR according to the number of oocytes retrieved in highly receptive patients under 40. (Polyzos et al;, 2018). This was confirmed by a retrospective study which concluded that the cumulative live birth rate significantly increased as the number of oocytes retrieved increased, up to 21-30 oocytes (Zhao et al., 2020). Recently, a prospective observational study concluded that CLBR increases according to the number of blastocytes formed, and so to the number of oocytes retrieved in the FAS (Papanikolaou E. et al., 2019).

Our own study concludes that for the FA group the cumulative live birth rate improves when up to 25 oocytes are retrieved. No advantage is gained by retrieving more. For the C group the figure is 20 oocytes. Noticeable also, in the freeze-all group, is the very poor performance when less than 6 oocytes are retrieved (20%) indicating a possible sub-group of patients for whom the indication of the freeze all is less advisable.

Equally of note, we retrieved more than 26 oocytes in 9% of the FA group whereas this was true only of 0.8% of patients in the control group.

### Strengths and limitations

The limitations of this study are, first and foremost, that it was carried out retrospectively and over a long period of time. Evolving methods and techniques, such as freezing and transfer methods, affected the results to some degree. Also, the power of this study is under 80% for some patient subgroups even with a large cohort. Another weak point is the heterogeneity of the freeze-all group. Concerning the obstetrical aspects, recent studies show a difference between fresh and frozen-thawed embryo transfers in neonatal and maternal outcomes which was not analysed in this study. (Ginström E. et al., 2019). A prospective randomised controlled trial is currently in progress and may reveal some answers. (Maheshwari A. et al., 2019). Finally, the cost effectiveness per ongoing pregnancy or per live birth was not analysed in this study. Even though, recent observational studies show that the FAS reduces the total cost compared to fresh cycles (Roque et al., 2015, Papaleo et al., 2017).

On the other hand, we should underline some strengths. This was a “real life” study and as such, reflected everyday patient realities, during normal daily practices, not made to conform to a particular methodology. In practice, we did not treat a selected group of women as we do in prospective randomised trials. Even if such studies give us precise evidence- based medicine, a “real life” study is a better guide to our daily work, where heterogeneous groups of patients are treated. This approach does not replace randomised controlled trials but comes as complementary information. The logistic regression allows to take into account the confounding factors and, to some extent, limits the bias of such retrospective studies.

The number of patients allows a comparison with a large control group, having undergone fresh, followed by remaining frozen-thawed embryo transfer. Also, our results analyse the cumulative live birth rate which allows more complete and interesting analyses than the live birth rate after a single transfer. As our control group included women with at least one frozen embryo in addition to the fresh one(s)it may be more appropriate to compare the results with a FA cohort rather than comparing it with patients having only a fresh transfer.

## Conclusion

Considering the cumulative life birth rate of the FA group and the C group (group of patients with fresh embryo transfer and possible additional frozen-thawed embryo transfer), the results are in favour of fresh embryo transfer, more efficient and faster retrospectively. Nevertheless, the CLBR in the FA group remains excellent and certainly good enough to maintain and support this approach for selected patients (Blockeel et al, 2019). Our results underline the advantages of retrieving large quantities of oocytes, especially for FA patients, and those of blastocyte vitrification, rather than slow freezing of cleaved stage embryos.

We confirm the high efficiency of FAS but underline the necessity to restrict this strategy to selected cases.

